# Recent speciation and adaptation to aridity in the ecologically diverse Pilbara region of Australia enabled the native tobaccos (*Nicotiana*; Solanaceae) to colonize all Australian deserts

**DOI:** 10.1101/2024.06.12.598428

**Authors:** Luiz A. Cauz-Santos, Rosabelle Samuel, Dominik Metschina, Maarten J.M. Christenhusz, Kingsley W. Dixon, John G. Conran, Ovidiu Paun, Mark W. Chase

**Author notes:** Author for correspondence: Luiz A. Cauz-Santos.

## Abstract

For the last six million years, the arid Australian Eremaean Zone (EZ) has been as dry as today. An accepted hypothesis, applied to arid regions worldwide, suggests that flora and fauna were more broadly distributed before aridification began. In Australia, this aridification process started around 20 million years ago (Mya), leading to gradual speciation processes via vicariance as the climate became increasingly arid. Here, we use genomic data to investigate the biogeography and timing of divergence of native allotetraploid tobaccos, *Nicotiana* section *Suaveolentes* (Solanaceae), which putatively entered the EZ 5 Mya. The original allotetraploid migrants from South America were adapted to mesic areas of Australia and putatively radiated recently in the EZ, including sandy dune fields (only 1.2 My old), after developing drought adaptations. Based on coalescent and maximum likelihood analyses designed to corroborate timing of the Australian radiation independently, arrival of *Nicotiana* section *Suaveolentes* on the continent occurred approximately 6 Mya, and ancestors of the Pilbara (Western Australian) lineages radiated there at the onset of extreme aridity 5 Mya by locally adapting to these various ancient, highly stable habitats. The Pilbara thus served as both a mesic refugium and cradle for adaptations to harsher conditions. This dual role is due to its high topographical diversity, providing microhabitats with varying moisture levels, and its proximity to the ocean, which buffers against extreme aridity. Consequently, species like *Nicotiana* have been able to survive in mesic refugia during arid periods and subsequently adapt to more arid conditions. These results demonstrate that initially poorly adapted plant groups can develop novel adaptations in situ, permitting extensive and rapid wide dispersal despite the highly variable and unpredictable extremes of heat and drought in the EZ.

## Introduction

The evolution of plant groups in the Eremaean Zone of Australia has been explained through three primary models: vicariance, pre-adapted immigration, and in situ adaptation. However, these models are not mutually exclusive and may operate concurrently or sequentially within different taxa. Vicariance, for instance, involves the gradual isolation and divergence of widespread taxa due to increasing aridity, as seen in *Eucalyptus* (Mytaceae) and *Acacia* (Fabaceae) (Martin, 2006; Byrne et al., 2008). In contrast, pre-adapted immigration, potentially (African or Asian) immigrant clades, supported by studies on *Triodia*, *Ptilotus* and other Amaranthaceae (Toon et al., 2015; Shepherd *et al*., 2004; Kadereit & Freitag, 2011; Hammer *et al*., 2021), suggests that some species arrived already equipped for arid conditions. Lastly, in situ adaptation, although less documented, involves the evolution of arid specialization within the Eremaean Zone itself. Australia has several of the oldest known pieces of the Earth’s crust (e.g., the Pilbara Craton, 3.8–2.7 Ga), but thanks to extensive erosion over such long timescales, these have a generally low-relief topography and few major physical barriers to dispersal, so it is possible that there is a third model: a group of organisms lacking xeric specializations could adapt in one region and then disperse over much of the Australian continent. These processes collectively contribute to the region’s complex biogeography and high levels of endemism.

Dispersal events, such as those facilitated by dry tornadoes (in Australia called willy-willies), have potentially played a crucial role in the diversification of *Nicotiana* in Australia. These events could enable the initial migration and subsequent wide distribution of *Nicotiana* species across the continent, contributing to the genetic isolation and diversification observed in the genus today. Unlike vicariance, which primarily isolates populations gradually, these dispersal events can rapidly introduce species to novel environments, creating new opportunities for speciation. However, the evolutionary history of *Nicotiana* section *Suaveolentes* has been the subject of different hypotheses. One hypothesis suggests that the origin of *N*. section *Suaveolentes* dates back to the early Miocene (ca. 20 Mya), with speciation occurring through a vicariant model of arid-adapted biota. This model, as described by Cracraft (1991) and Ladiges et al. (2011), proposes that species were progressively isolated as aridification moved from north to south, forming organized "tracks" of species distributions. Following these primary vicariance-driven distributions, dispersal events produced a few widespread species. The alternative hypothesis points that *Nicotiana* only reached Australia around 4–6 Mya, as suggested by Mummenhoff and Franzke (2007) and Schiavinato et al. (2020), with a more recent radiation occurring in the Eremaean Zone (Clarkson et al., 2017; Dodsworth et al., 2021). In this scenario, dispersal to the arid interior would have been the predominant mechanism of diversification, with vicariance playing a more secondary localized role.

The Australian distribution of *Nicotiana* section *Suaveolentes* spans the continent, except for Tasmania, which putatively makes them ideal for revealing general factors contributing to arid zone speciation. Furthermore, elucidation of the phylogeographic history for such widespread groups can be instrumental to understand the major drivers of individual species distributions. The Australian species are primarily found in the tropics, subtropics and warm temperate regions and largely absent from cool temperate regions (Ladiges *et al*., 2011). There was putatively rapid initial speciation producing a few species in more mesic areas in northern and eastern Australia, followed by multiple radiations in the EZ (Ladiges *et al.,* 2011, Chase *et al*., 2018, 2022a). The current understanding of the biogeography, phylogenetics and chromosome numbers/genome sizes of *N.* section *Suaveolentes* (Table 1; Chase *et al*., 2002b) are consistent with the ancestral distribution of the genus in Australia being confined to these northern and eastern parts, where all species with higher chromosome numbers (*n* = 22–24) and mostly larger genomes now occur (Fig. 1). As new species appeared and chromosome numbers and generally genome sizes decreased (Chase *et al*., 2002b), they expanded to the dry interior of central and southern Australia, where they are now highly diverse. Although several EZ species are specialists of more mesic niches, growing in the shade of trees or rock outcrops, many others inhabit exposed, extremely arid sites (Fig. 1), unexpected for such thin-leaved plants with no obvious adaptations for these extreme habitats. However, their rapid lifecycle completion following rainfall could sometimes allow them to avoid the harshest conditions.

**Fig. 1.**
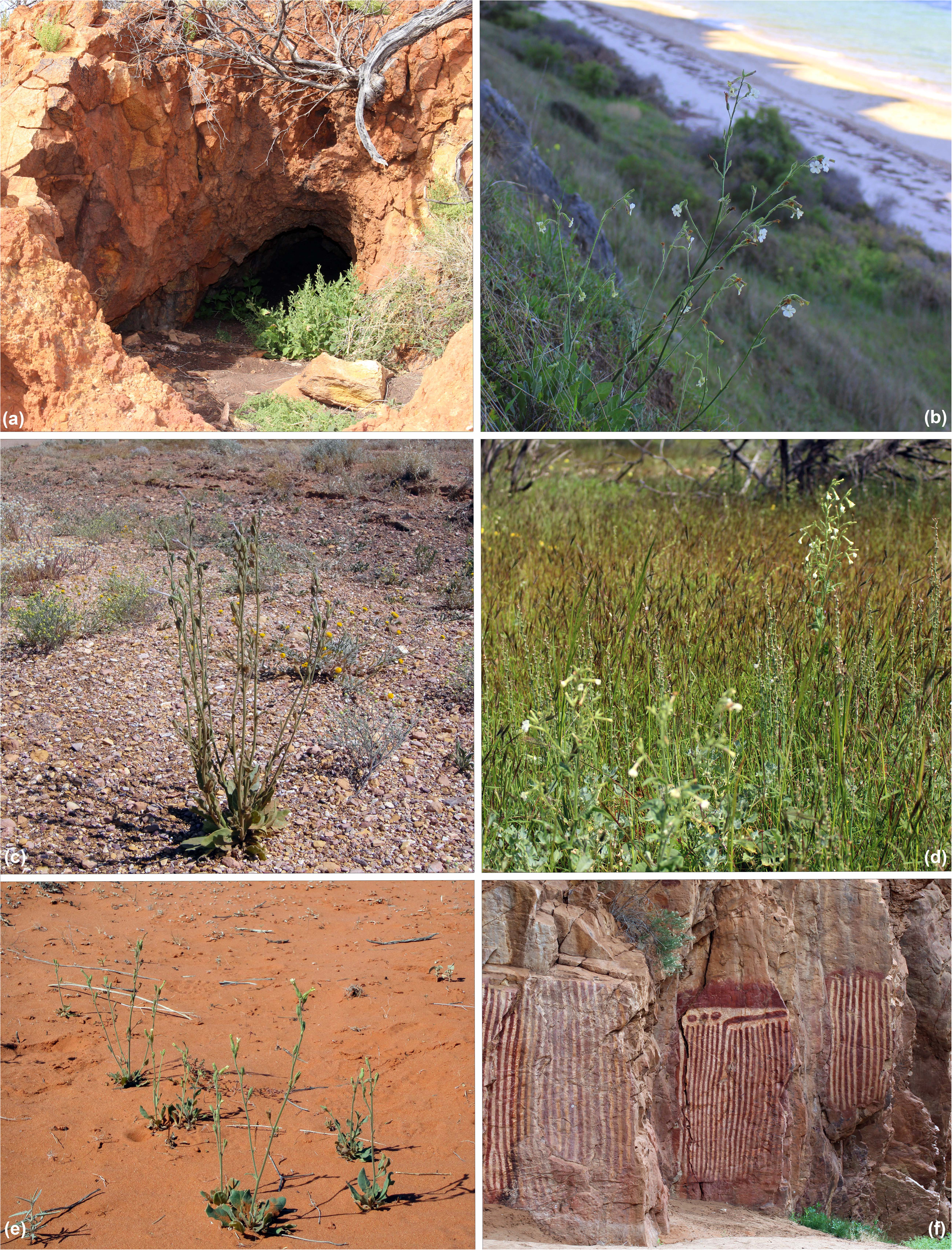
Habitat diversity, mesic and arid, among species of *Nicotiana* section *Suaveolentes* in the Australian Eremaean Zone. (a) Mesic site for *N. cavicola* near the entrance to a cave near Meekathara (Western Australia). (b) Mesic site for *N. maritima* on the sea cliffs near St. Vincent, York Peninsula (South Australia). (c) Arid site for *N. simulans* on the gibber plains south of Ooodnadatta (South Australia). (d) Mesic site for *N. insecticida* in a mulga woodland (*Acacia aneura* species complex) near Carbla (Western Australia). (e) Arid site for *N. velutina* in a fossilised sand dune near Coober Peddy (South Australia). (f) Mesic site for *N. insecticida* in the Anthwerrke Gap, East MacDonnell Ranges (Northern Territory).

**Table 1.** Biogeographic stochastic mapping (BSM) counts for *N*. section *Suaveolentes* using the Australian drainage regions as operational areas and the DIVALIKE+J model selected in BioGeoBEARS. Mean values (mean) and standard deviations (stdev) are event counts of 100 BSMs.

| Mode | Type | mean | stdev | % |
| --- | --- | --- | --- | --- |
| Within area speciation | Sympatry - narrow | 22.98 | 1.59 | 36.47 |
| Dispersal | Jump-dispersal | 38.23 | 1.69 | 60.68 |
| Vicariance |  | 1.79 | 0.46 | 2.84 |
| Total events |  | 63 |  |  |

Here, we have used multispecies coalescent methods on genome-wide single nuclear polymorphisms (SNPs) combined with biogeographical and novel molecular clock analyses to evaluate these two competing hypotheses that the radiation of *N.* sect *Suaveolentes* in the EZ started either as early as 20 Mya or as late as 6 Mya. The former would coincide with the early stages of aridification in Australia, supporting the vicariance model where species gradually became isolated and diversified as the climate changed (Ladiges *et al.,* 2011), whereas if the latter is true then dispersal was the preponderant mechanism and diversification took place long after the EZ became dry. Inadequate, fine-scale sampling in such remote areas across the whole continent has limited previous phylogeographic studies, and existing phylogenetic analyses typically have not sampled species thoroughly either through lack of sufficient collections or due to the existence of many undescribed “cryptic” species. After ten years of fieldwork, we have assembled an unprecedented collection of accessions and nearly trebled the number of species known in this group (Chase *et al*., 2021, 2022, 2023). We have also extended our taxonomic coverage by recovering additional taxa/accessions from viable seeds on herbarium specimens up to 25 years old. We first constructed a species tree for *N.* sect *Suaveolentes* and implemented dating of the speciation events to corroborate previous molecule-based age estimates (Clarkson *et al*., 2017; Schiavinato *et al*., 2020; Dodsworth *et al*., 2021). We then carried out biogeographical inference to test the relative contribution of dispersal and vicariance to the overall distribution of the species. Our primary goal is to identify where and when the initial lineages confined to the wetter parts of the Australian continent became adapted to xeric conditions, providing insights to how they diversified into the myriad, relatively unoccupied niches hypothesized to be typical of arid zones.

## Material and Methods

### Plant material

Sampling of 273 accessions from 58 species of *N.* section *Suaveolentes* (Table S1), extended the recent studies of Cauz *et al*. (2022), which focused on *N. benthamiana*, and Chase *et al*. (2022), which examined genome size and chromosome number evolution in this section of the genus. Most samples were collected by Chase and Christenhusz in the wild and are vouchered in major Australian Herbaria (AD, BRIS, CANB, CNS, DNA, NSW, NT and PERTH, the standard acronyms for these collections). Other accessions were added by germinating viable seeds removed from herbarium specimens. From more than 600 accessions, including roughly 100 from herbarium-stored seeds, we selected the above subset of 278 samples that it comprised all putative species, including several that are undescribed and designated here as sp. nov. with a location where we collected these taxa (e.g., *N. sp. nov.* Coondiner). The maps were generated in QGIS v. 3.20.3 (QGIS Development Team, 2021); the map layer with the vectors from the drainage and rivers divisions were obtained from the Australian Government Bureau of Meteorology (http://www.bom.gov.au/water/geofabric/). The provenance data (latitude, longitude) of the accessions were obtained from the Australasian virtual herbarium (https://avh.chah.org.au) and our personal data gathered during the field collections in Australia.

### Collecting and import permits

The following collecting permits, which cover these accessions, were issued to MWC and MJMC: Western Australia SW017148, CE006044, Northern Territory 58658, and Queensland PTU-18001061. Removal of seeds from herbarium specimens was approved by the curators/collections managers of the following herbaria: AD, BRI, CANB, NSW, NT and PERTH. All seeds imported into the UK followed published guidelines; plants were grown at the Royal Botanic Gardens, Kew, under DEFRA PHL2149/194627/5NIRU CERT:106-2019; HMRC TARIFF CODE: 0601209090. No material collected by us or in our possession will be exploited for commercial purposes without involvement of the Australian and Aboriginal authorities, as required by the collecting/export permits.

### DNA isolation, library preparation and sequencing

From ca. 20 mg of silica-dried leaf tissue, we extracted DNA with the cetyltrimethylammonium bromide (CTAB) procedure (Doyle, 1990), following a 20 min ice-cold sorbitol buffer treatment (100 mM tris-HCl, 5mM EDTA, 0.35 M sorbitol, pH 8.0). Then, we used 2.5 μl of RNase A (Thermo Fischer, USA) for 30 min at 37 °C and purified the extracted DNA using the NucleoSpin gDNA clean-up Kit (Machery-Nagel, Germany), according to the manufacturer’s instructions.

DNA samples were first single digested with the PstI restriction enzyme in advance of library preparations. Although PstI activity is not affected by CG methylation, it is sensitive to that at CHG sites (H stands for any nucleotide apart from G), a type of methylation frequently found around plant transposable elements (e.g., Domb *et al*., 2020), which are known to have paralogy issues and low phylogenetic signal. The effects of any methylation variation have been mitigated by filtering for missing data. Libraries were prepared following Paun *et al*. (2016), as modified by Cauz *et al*. (2022) and Chase *et al*. (2022). Processing in batches was carried out using index barcodes distinct from one another by at least three bases. Sequencing was performed at the VBCF NGS Unit (www.vbcf.ac.at/ngs) on an Illumina HiSeq 2500 with 125 bp paired-end reads.

### SNP calling and phylogenomic analysis

The BamIndexDecoder v.1.03 (included in Picard Illumina2Bam package, available from http://gq1.github.io/illumina2bam/) was used first to process the RADseq data and demultiplex via the index barcodes in sublibraries. Subsequently, demultiplexing of individuals via their inline barcodes was conducted in process_radtags from Stacks v.1.47 (Catchen *et al*., 2013), together with removal of reads containing uncalled bases or with low quality-scores.

The reference genome of *N. benthamiana* v.2.6.1 (Bombarely *et al*., 2012, available from https://solgenomics.net/organism/Nicotiana_benthamiana/genome), a member of *N.* section *Suaveolentes*, was used on individual read mappings in BWA MEM v. 0.7.17 (Li and Durbin, 2009) using and applying the –M option to flag shorter splits hit as secondary. We also checked for biases potentially driven by phylogenetic relatedness to the reference individual. After alignment, the sam file was sorted by reference coordinates, and read groups were added using Picard Toolkit v.2.27 (available from http://broadinstitute.github.io/picard/). We used the Genome Analysis Toolkit (GATK) v.3.8 (McKenna *et al*., 2010) to improve alignment quality around indels, thinning the data to a maximum of 100,000 reads per interval.

GATK was used to call variants following the best-practice DNAseq recommendations. First, we inferred genotypes via HaplotypeCaller and GVCF mode for individual samples and subsequently processed all individual intermediate GVCF in a joint genotyping analysis in the GenotypeGVCFs module. The raw vcf file was first processed in VCFtools v.0.1.15 (Danecek *et al*., 2011) and retained only variants presents in at least 50% of individuals. We then used the VariantFiltration GATK module with the following criteria: (1) depth of coverage (DP) < 500; (2) variant confidence (QUAL) < 30.00; (3) variant confidence divided by the unfiltered depth (QD) < 2; (4) Phred-scaled P-value for the Fisher’s exact test to detect strand bias (FS) > 60; (5) a root mean square of mapping quality across all samples (MQ) < 40; (6) u-based z-approximation from the rank sum test for mapping qualities (ReadPosRankSum) < -8.0; and (7) u-based z-approximation from the rank sum test for the distance from the end of the reads with the alternate allele (MQRankSum) < - 12.5.

The variant calling and initial filtering steps in GATK produced 7,606,626 variable sites, but we retained only SNPs with a minor allele frequency ≥ 0.008 (i.e., present in at least four haplotypes), an average depth above 20 and 20% maximum missing data. We also filtered the data using the populations pipeline in Stacks to retain only variable positions with a maximum observed heterozygosity of 0.65, thus avoiding the use of pooled paralogs in further analyses.

To investigate phylogenetic relationships among the species of *N.* section *Suaveolentes*, we first converted the final filtered vcf to a PHYLIP file using PGDspider v.2.1.1.0 (Lischer & Excoffier, 2012). We removed invariant sites with the script ascbias.py (https://github.com/btmartin721/raxml_ascbias). A RAxML v.8.2.12 (Stamatakis, 2014) used the remaining 170,552 SNPs with the recommended ascertainment bias correction (Lewis, 2001). The phylogenetic tree was inferred under the GTRCAT model of nucleotide substitution with a search for the best-scoring ML tree and 1,000 rapid bootstrap replicates. We assigned *N. africana* as the outgroup because it was well-supported sister to the rest of *N.* section *Suaveolentes* in all previous studies (Chase *et al*., 2003; Clarkson *et al*., 2004, 2010, 2017; Marks *et al*., 2011; Kelly *et al*., 2013). All species of *N.* section *Suaveolentes* are allotetraploids, including *N. africana* and *N. benthamiana*, so our analyses do not mix diploid and polyploid taxa. Finally, we visualized and annotated the best tree in R, using ape v.5.3 (Paradis & Schliep, 2018), biostrings (Pagès *et al*., 2020), ggplot2 (Wickham, 2016), ggtree (Yu *et al*., 2017) and treeio (Wang *et al*., 2020).

### Coalescent-based species tree and divergence time estimation

To construct a species tree for a reduced and representative matrix of *N.* section *Suaveolentes*, we first inferred the relatedness between accessions looking for evidence of introgression, which would interfere with species-tree inference. This exercise was conducted on a reduced matrix of 22 species representing the major clades in *N.* section *Suaveolentes*. After calculating genotype likelihoods in ANGSD v.0.930 (Korneliussen *et al*., 2014) and retaining only variants with a minor allele shared by at least two individuals, we applied a minimum base mapping quality of 20, SNP calling confidence of p<1e^-6^, and presence in at least 70% of individuals. The major and minor allele frequencies were estimated using the GATK-based genotype likelihood model, and our final dataset resulted in 3,201,820 variable positions. For the inference of coancestry, we used our estimated genotype likelihoods to obtain a covariance matrix using PCangsd (Meisner & Albrechtsen, 2018), and plotted our data using the heatmaps.2 function from GPLOTS v.3.0.1.1 (Warnes *et al*., 2020).

To construct the coalescent species tree, we first used the filtered vcf file to prepare a smaller dataset in VCFtools v.0.1.15 selecting only unlinked biallelic SNPs (> 10,000 bp apart on a contig) and removing missing data at each locus. This procedure produced a matrix of 2,400 unlinked single nucleotide polymorphisms (SNPs) for 36 accessions, representing 18 species (two per species) that broadly cover the phylogenetic diversity of *N.* section *Suaveolentes*. We converted the vcf file containing unlinked SNPs to PHYLIP and then NEXUS format using PGDSpider v.2.1.1.0, and finally we created the input XML files in BEAUti v.2.4.8 (Bouckaert *et al*., 2014).

We constructed the coalescent species tree using SNAPP v.1.2.5 with a chain length of 10 million and saving a tree every 1,000th generation. We monitored the convergence of the run based on the ESS values from the log-file with Tracer v.1.6 (Rambaut *et al*., 2018). We removed the initial 10% of trees as burn in, visualized the SNAPP trees as a cloudogram using Densitree v.2.2.6 (Bouckaert & Heled, 2018) and produced the posterior probabilities for each clade with Treeannotator v.1.8.3 (Drummond *et al*., 2012). To calibrate the species tree, we used 5e^-09^ as the rate of substitution per site per generation (Schiavinato *et al*., 2020) and one year as generation time (i.e., these plants only rarely live more than one season in nature). We estimated divergence times by rescaling the results using the total length of investigated sites for the loci included and total number of polymorphic sites across their length.

### Ancestral range estimation

For biogeographic inference, we calibrated a RAxML tree using TreePL (Smith & O’Meara, 2012), which produces a dated tree using a penalized likelihood approach, and minimum and maximum ages to constrain the tree. The minimum and maximum ages used for TreePL dating were based on the divergence times estimated in our SNAPP species tree. The dated tree with node ages was visualized using FigTree v1.3.1 (http://tree.bio.ed.ac.uk). We obtained the confidence intervals for the node ages and the maximum clade credibility (MCC) tree summarizing the RAxML bootstrap replicates with Treeannotator v.1.8.3 (Drummond *et al*., 2012) (Fig. S1).

We explored the biogeographic history of *N.* section *Suaveolentes* first by comparing models in BioGeoBEARS, including DEC, BAYAREALIKE and DIVA plus using the additional free parameter “j” in each model, which accounts for jump dispersal/founder-event speciation. According to the Akaike information criterion (AIC), the best-fit model was DIVA+J (Table S2). We then performed the ancestral range estimation using BioGeoBEARS testing two models of distribution. The first one considered drainage divisions of Australia as operational areas: A, Africa; B, Pacific, C, Carpentaria Coast; D, Tanami-Timor Sea Coast; E, north-western Plateau; F, Pilbara-Gascoyne; G, Southwest Coast; H, Southwest Plateau; I, Lake Eyre Basin; J,LMurray-Darling Basin; K, Northeast Coast (Queensland); L,LSoutheast Coast (NSW); M,LSoutheast Coast (Victoria; N, South Australian Gulf. We chose this model because we had noticed in a previous study (Cauz-Santos *et al*., 2022) that species distributions seemed to conform to river drainage basins. Rather than acting as barriers to gene flow, it is more likely that the drainages serve as conduits, facilitating gene flow through the dry tornadoes that typically travel across relatively flat landscapes and dissipate upon encountering uneven terrain. Consequently, dispersal mostly occurring within a drainage system might make sense in this setting, an aspect that we are planning to investigate further.

The second model split *N.* section *Suaveolentes* into the operational areas from Ladiges *et al*. (2011): A, Africa, Namibia; B, Pacific; C, eastern Australia; D, south-eastern Australia; E, south-eastern Interzone; F, north-eastern Interzone 3; G, Adelaide/Eyre; H, south-western Interzone; I, Pilbara; J, north-western Australia; K, western Desert; L, northern Desert; M, eastern Desert; N, Nullarbor; O, central Australia. These geographical areas were defined by Ladiges *et al*. (2011) based on the distribution of narrow-range endemic species, Australian bioregions recognized by other authors (Burbidge, 1960; Cracraft, 1991; Crisp *et al*., 1995) and the Interim Biogeographic Regionalisation of Australia (IBRA) v.6.1 (http://www.environment.gov.au). The BioGeoBEARS model testing for this distribution also resulted in the DIVA+J as best fit AIC model (Table S3). In both models, the species distributions (presence/absence) included a maximum of three areas per species except for one widespread species, *N. velutina*, although we now believe that this species is much more restricted than previously thought due to discovery that this species concept as previously defined includes two previously unrecognized species, considerably reducing its distribution. However, all three form an exclusively related species complex, so in terms of these analyses, this is an acceptable assumption. The accessions included in our analyses are from only one of the revised concepts in the *N. velutina* complex (Cauz-Santos, Metschina & Chase, unpubl.).

## Results

### Phylogenetic analysis of *Nicotiana* section *Suaveolentes*

We obtained an average of 3,053,843 paired-end reads for the 273 accessions used in this study. The filtered reads mapped onto the *N. benthamiana* reference genome at a high rate (an average of 95.75%), with a final average coverage across samples of 9.4. The mapped reads were then used for variant calling, which after filtering resulted in a total of 240,871 SNPs for a minimum of 80% of individuals. The results shown here expanded the matrix published in Chase *et al*. (2022) to include many more of the new species. These results were also used to illustrate the positions of species being described as new (Chase *et al*., 2023, in press), but the methods and results of this analysis are here published for the first time. Phylogenetic trees of *N.* section *Suaveolentes* (Fig. 2) provide a framework for the other studies conducted for this paper, but they are not a primary focus here and have not been used previously for these purposes.

**Fig. 2.**
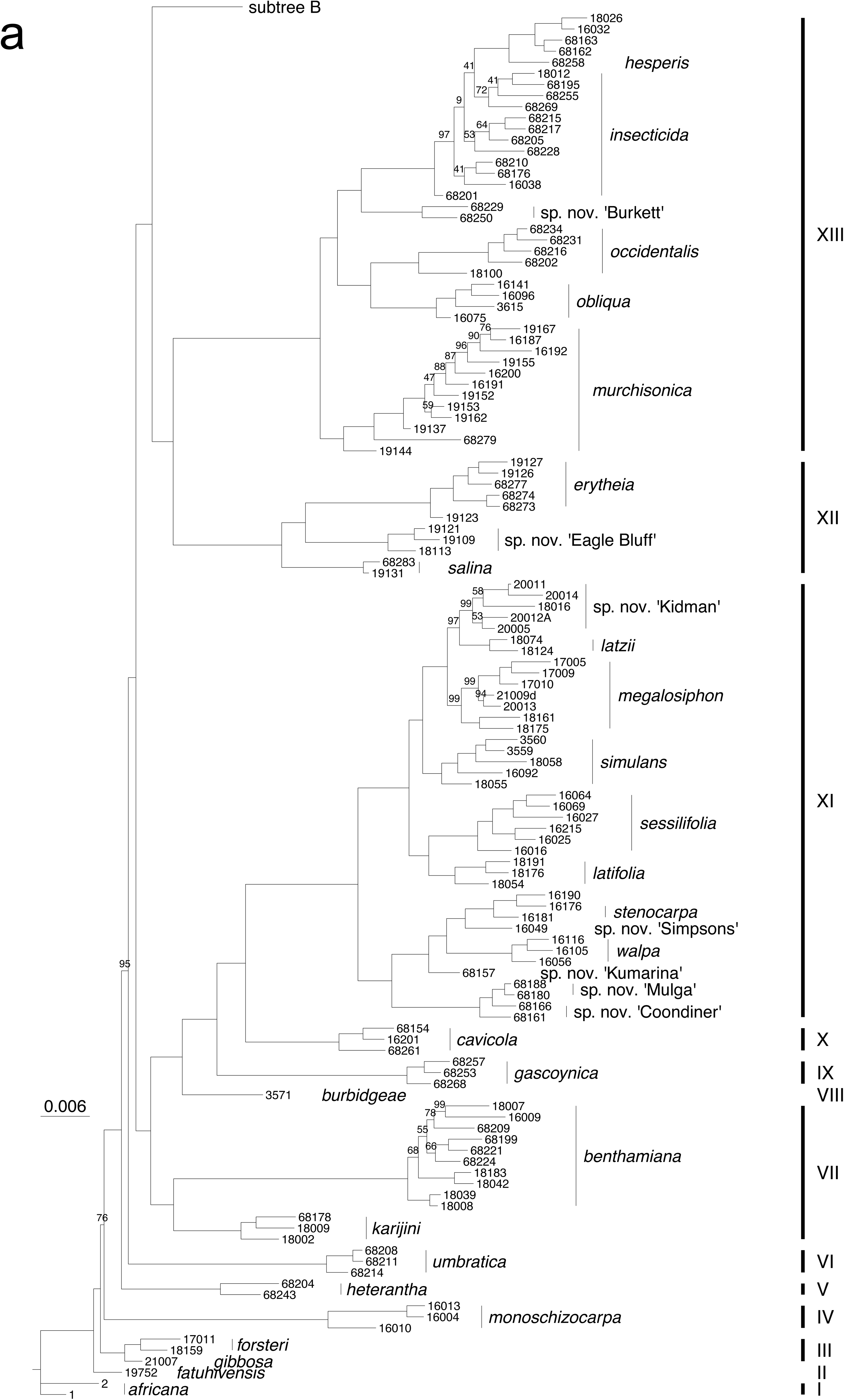
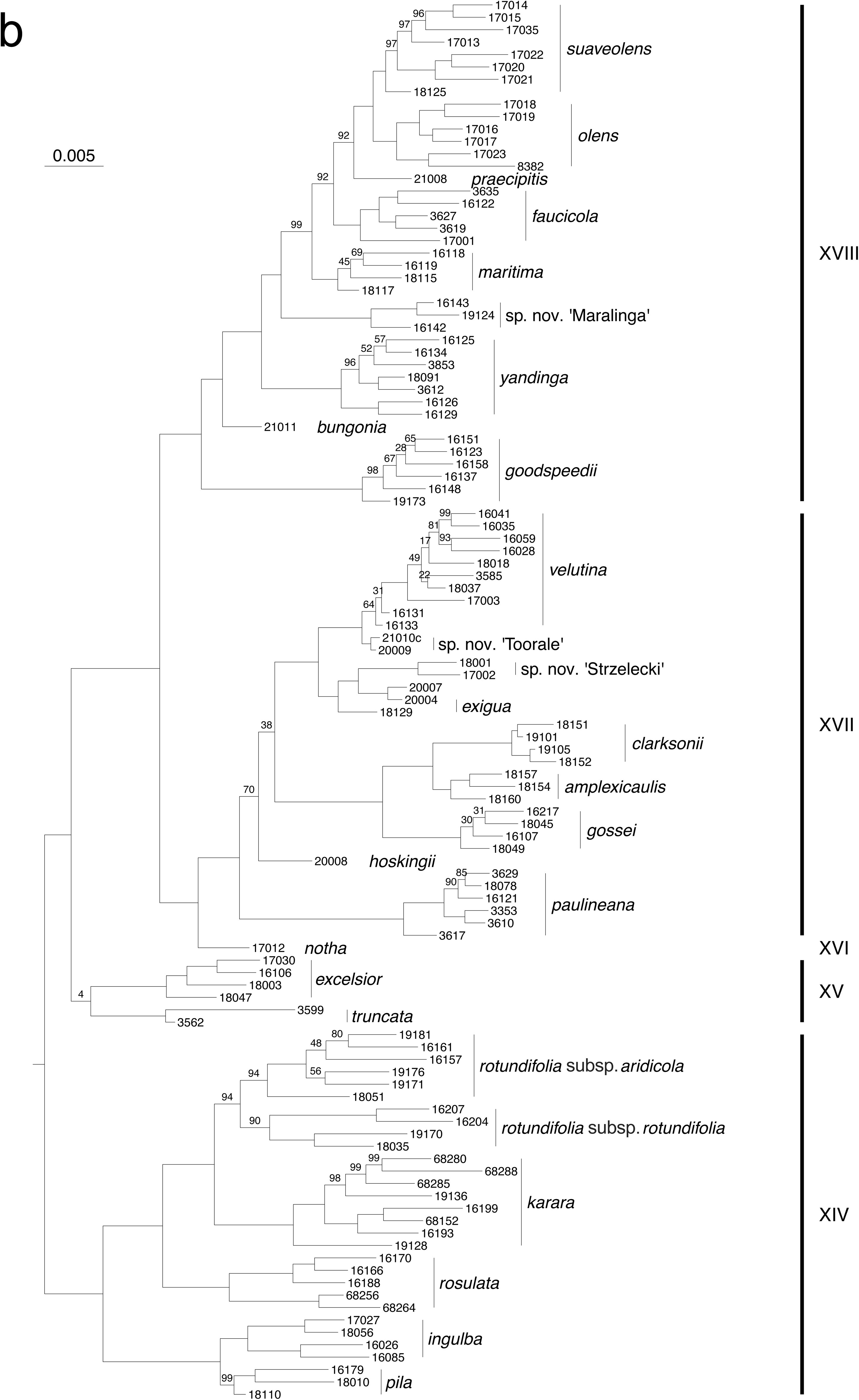
RAxML phylogenetic tree from *N.* section *Suaveolentes*, subtrees (a) and (b), based on 240,871 SNPs. The numbers are bootstrap percentages; branches without numbers received 100.

The RAxML maximum likelihood (ML) tree was in general highly supported (most nodes with bootstrap percentage, BP, >90; Fig. 2). For several species, we included multiple accessions that formed unique and well-supported groups. In total, the tree comprised 18 major clades (numbered as Roman numerals I–XVIII), with *N. africana* (I) outgroup to the rest of *N.* section *Suaveolentes* (Clarkson *et al*., 2011). The basal node comprises *N. fatuhivensis* (II) sister to *N. gibbosa*+*N. forsteri* (III), but with low support (BP 76; the only BP less than 90 along the spine of the ML tree) relative to the position of *N. monoschizocarpa* (IV). *Nicotiana heterantha* (V) and *N. umbratica* (VI) are then successively sister to the rest, which split into six geographically widespread clades (VII, VIII–XI, X/XI, XII/XIII, XIV, XV–XVIII).

### Species tree and divergence times

The coancestry heatmap (Fig. 3) shows the clear relatedness among accessions of *N. monoschizocarpa*, which is one of the species at the basal node of the *N*. section *Suaveolentes*, but this pattern is not evident in the other two species near the basal nodes, *N. africana* and *N. forsteri*. *Nicotiana africana* grouped within the *N. forsteri* accessions with high coancestry, and additionally, one of the *N. africana* accessions exhibited high coancestry with the species of the *N. benthamiana* complex. Considering these patterns of introgression and as only two accessions of *N. africana* have been available to us (a minimum of two accessions per species is required to produce the species tree), we decided to remove *N. forsteri* and species of the *N. benthamiana* complex from the species tree inference. The highest coancestry is exhibited by the most recent species groups, *N. gossei*/*N. velutina*, *N. truncata*/*N. excelsior* and *N. maritima/N. suaveolens*. Introgression is not a general phenomenon in *N*. section *Suaveolentes*, but it does seem to be a factor for a few species pairs.

**Fig. 3.**
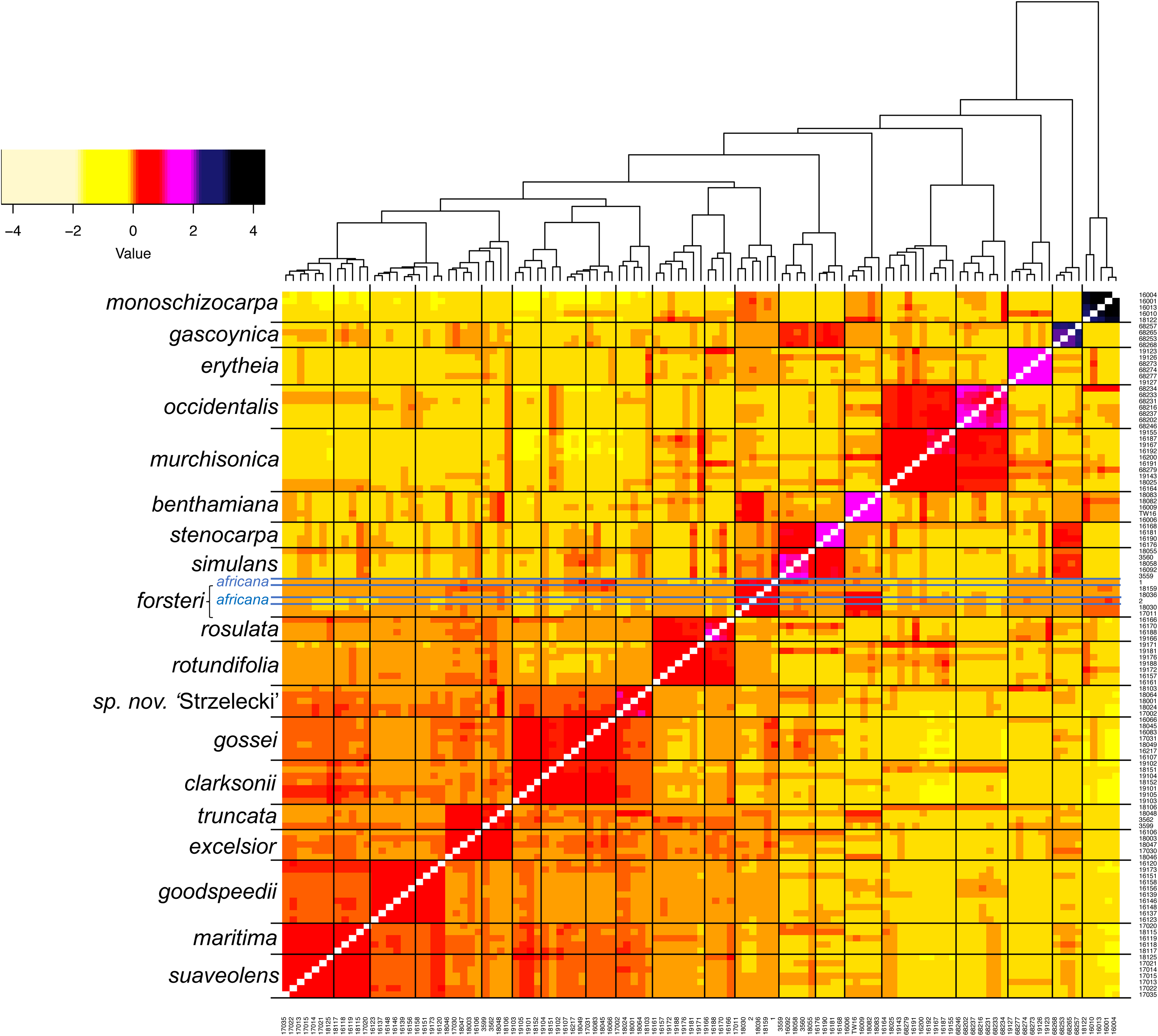
Coancestry heatmap from the pairwise relatedness between accessions from *N*. section *Suaveolentes*. The darker colours indicate higher relatedness; the estimates of relationship of individuals to themselves are not shown. The blue lines specify the accessions of *N. africana*. High levels of coancestry among recently diverged species (e.g., *N. goodspeedii/N. maritima/N. suaveolens*) are to be expected.

The coalescent species tree for a reduced dataset comprised representative species from all major clades in *N*. section *Suaveolentes*. We obtained one topology in the SNAPP analysis with strong support representing 99% of the posterior density distribution (Fig. 4). The ESS values were used to monitor the convergence of the analysis, from which we obtained values higher than 200 in each parameter. The species tree places *N. africana* as sister to the rest, consistent with its outgroup position in the ML results (above) followed by *N. monoschizocarpa*. Subsequently we observed three clades, one comprising (*N. occidentalis*+*N*. *murchisonica*)+*N*. *hesperis* (clade XII/XIII in the ML tree), a second with (*N. gascoynica*+*N. simulans*)+*N. stenocarpa* (clade VII–XI in the ML tree) and a major clade (*N. rotundifolia* to *N. gossei*) comprising species from the two remaining major clades in the ML tree (XIV, XIV–XVIII).

**Fig. 4.**
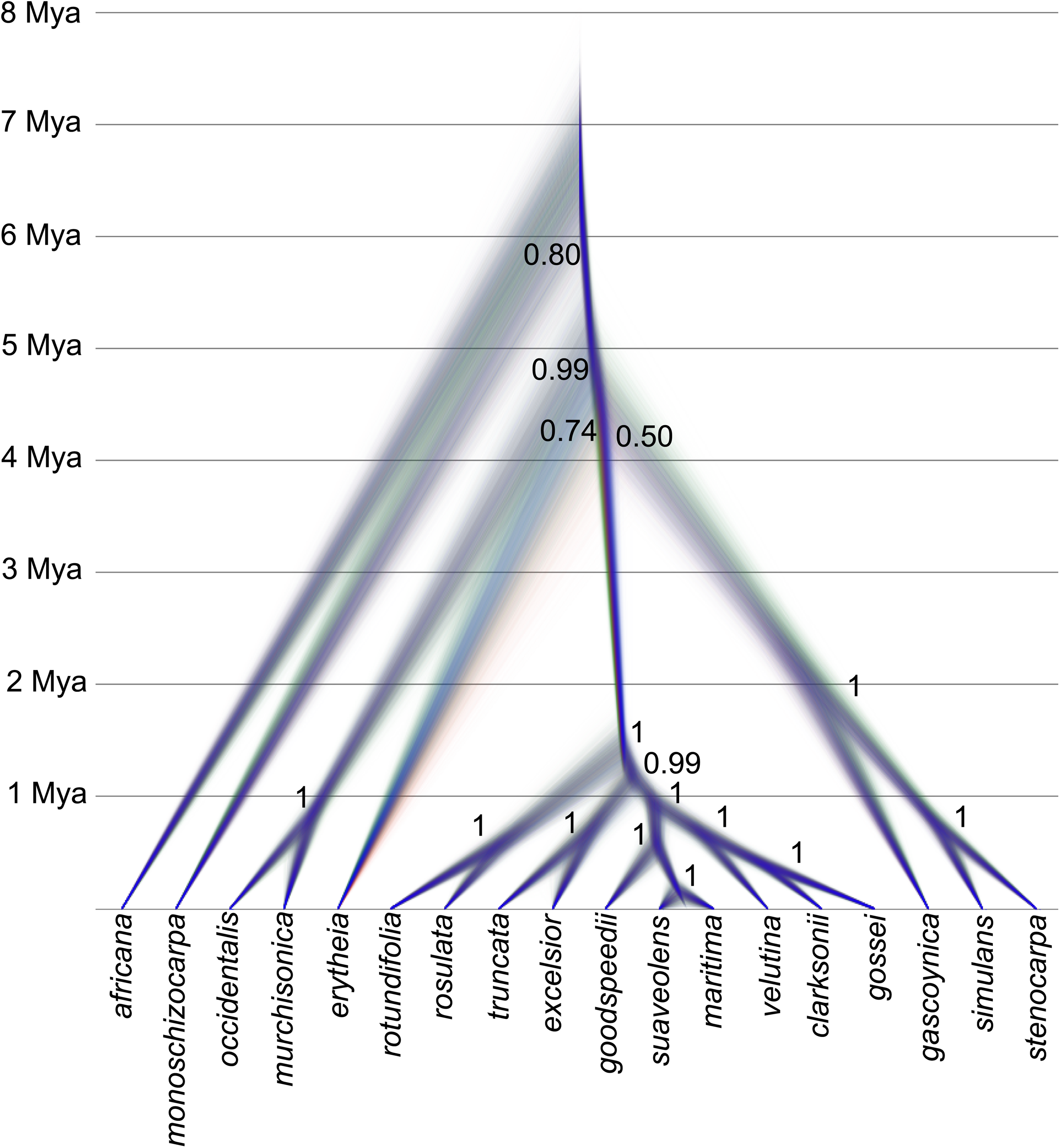
Cloudogram from species trees generated for *Nicotiana section Suaveolentes*, scaled to divergence times. The species tree reveals three of the main radiations in the section, the *N. occidentalis*, *N. gascoynica* and *N. suaveolens* clades, the last being the most recent radiation (ca. 1 Mya) comprising the species occurring in central and southern Australia.

In the divergence time estimation, *N. africana* diverged from the remainder of the section around 6.5 Mya, in the late Miocene. Among the Australian species, the first split, *N. monoschizocarpa* from the rest, was estimated at 6 Mya, followed by clades XI/XII and VI– XI clades, at 5 and 4 Mya, respectively. Finally, the largest clade, XIV–XVIII, diversified in the last 1 My.

### Biogeographic history of *Nicotiana* section *Suaveolentes*

For the biogeographic analysis, we first evaluated the best model for our dataset including the possibility of founder-event speciation or jump dispersal (adding the j parameter in BioGeoBEARS). According to AIC values, the DIVALIKE+J model was best, accounting for 79% of the predictive power found in all tested models. This model allows for the possibility of anagenetic (dispersal and extinction) and cladogenetic events (vicariance), and the J parameter added to the model the possibility of founder events. Considering our favored distribution model (Australian river drainages) and the fact that most species do not occur in more than one drainage basin, selection of DIVALIKE+J has biogeographical support.

The ancestral area reconstruction (DIVALIKE+J and Australian river drainages as divisions for the distribution species in *N.* section *Suaveolentes*) resulted in a total of 63 biogeographic events (Table 1, Fig. 5). The results show a combination of anagenetic and cladogenetic events playing a role in the distribution of this group, with founder events (60.6%) being the main source of speciation, followed by within-area speciation events (36.4%) and a small proportion of vicariance (3.6%). In the biogeographic tree, the arrival of the ancestor of the *N*. section *Suaveolentes* in Australia occurred around 5.2 Mya. Even if the ancestral range of the species at the basal nodes is unclear (*N*. *monoschizocarpa*, *N. gibbosa* and *N. forsteri*), the common ancestor of the rest expanded its distribution around 5 Mya to the Pilbara region (F), in which several within-area speciation events occurred, resulting in this region becoming a *Nicotiana* biodiversity hotspot. The Pilbara region (F) as referred in this study encompasses the Pilbara Craton and adjacent basins of the Gascoyne, Wooramel, and Murchison Rivers in north-western Western Australia (north of latitude 25°00′ S and west of longitude 121°30′ E; Pepper *et al*., 2013) (Fig. 5). This area is larger than the Pilbara Bioregion of the IBRA classification, which only covers a portion of region (F), and is characterized by its unique geological formations, including the ancient Pilbara Craton, and features a variety of landscapes such as mountain ranges, coastal plains, and arid desert areas. From the Pilbara, a series of dispersals to the other parts of the EZ occurred, the last of which was to central and southern Australia. The second model using the areas as defined in Ladiges *et al*. (2011) also resulted in the Pilbara (region I) being colonized by the species of *N.* section *Suaveolentes* around 5 Mya, with a series of dispersal events from there, leading to the current distribution of this section in central and southern Australia (Table S4, Fig. S2).

**Fig. 5.**
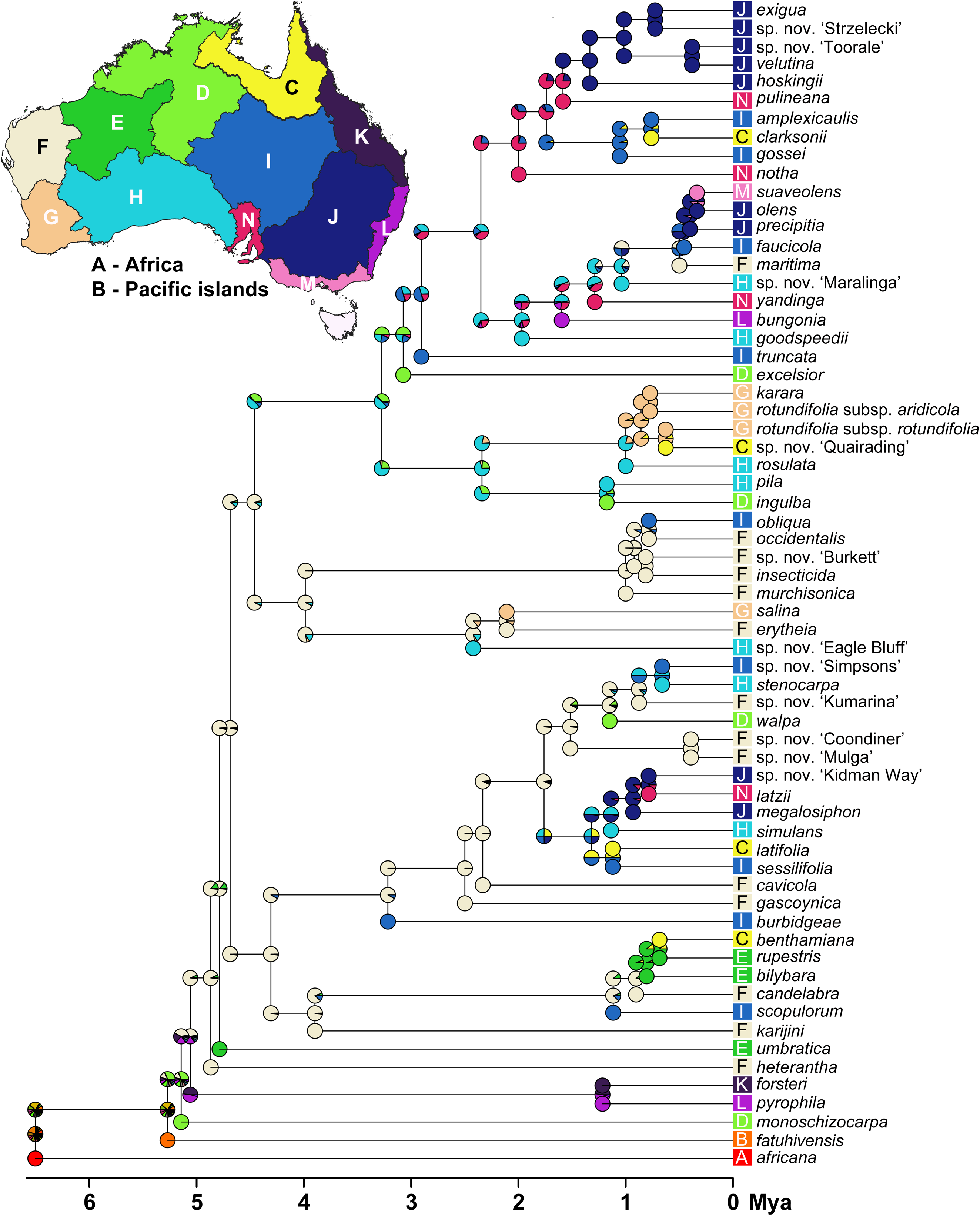
Biogeographic history of *N.* section *Suaveolentes*. Ancestral area reconstruction using the drainage areas as a model and showing greater Pilbara region (region F/light green) as the ancestral range for the species adapted to the Eremaean Zone (ca. 5 Mya). The map in the upper part of the figure represents the biogeography model used for distributions of the species in *N.* section *Suaveolentes*. The map shows the Australian drainages employed here to evaluate dispersal versus vicariance during diversification of *N.* section *Suaveolentes*.

## Discussion

### Timing of diversification in the Pilbara region and dispersal to the rest of the Eremaean Zone

The evolutionary processes shaping arid-adapted biota are complex and multifactorial, involving in many cases processes as vicariance, isolating populations due to climatic or geological changes, long-distance dispersal, enabling colonization of new habitats, and in situ speciation, driven by microhabitat differentiation and local environmental pressures. These mechanisms interact dynamically, creating genetic diversity and endemism. This complexity, influenced by the harsh and fluctuating conditions of arid environments, results in unique evolutionary trajectories. Other studies on arid-adapted biota have already demonstrated the complexity of evolutionary models in these environments (Cracraft, 1991; Byrne et al., 2008; Ladiges et al., 2011).

Our study contributes to this understanding by showing that *N.* section *Suaveolentes* first occupied mesic regions of Australia around 6 Mya. Subsequently, these species adapted to the arid conditions of the Pilbara region around 5 Mya, leading to their widespread distribution across the Eremaean Zone. This pattern of adaptation and dispersal is significant within the context of the Australian flora, and while highlights a specific example within *N.* section *Suaveolentes*, it is important to recognize that similar processes may have occurred in other plant groups, and further studies may reveal additional examples of such evolutionary dynamics. Molecular clock methods using multiple calibrations based on species pairs on oceanic islands/mainland (Clarkson *et al*., 2005), secondary ages from other analyses (Clarkson *et al*., 2017), and various methods of estimating rates of molecular divergence (Mummenhoff & Frankze 2007; Schiavinato *et al*., 2021; this paper) produce relatively consistent age estimates for formation of *N.* section *Suaveolentes,* ca. 5–7 Mya. Ladiges *et al*. (2011) disparaged the estimate of Mummenhoff & Frankze (2007) by citing other molecular clock studies (more generally on Australian angiosperms) that have provided highly divergent timings, but those for *Nicotiana* using these several approaches thus far have been relatively consistent for *N.* section *Suaveolentes*.

We found dispersal to be the predominant mode of evolution for *N.* section *Suaveolentes* throughout the EZ with some local examples of *in situ* speciation and vicariance being secondary features. Diversification in each major clade appears to be recent and ongoing, including adaptation to sandy dune fields (e. g., *N. pila*, *N. inglba*, *N. latzii*, *N. velutina*), which only appeared c. 1.2 Mya (Fujioka *et al*., 2009), concurrent with lower frequency/higher amplitude glacial cycles worldwide c. 0.8–1.2 Mya (Clark *et al*., 1999). The recentness of speciation accounts for the cryptic nature of many recently described *Nicotiana* species (Chase *et al*., 2023).

However, diversification has not been limited to recent times, and our phylogeny (Figure 5) reveals significant speciation events that occurred over a prolonged period and across multiple regions (for example in areas D and H). This pattern highlights a complex evolutionary history where both recent diversification (approximately 1 Mya) and older events have played crucial roles in shaping the current diversity of *N.* section *Suaveolentes*. The presence of these older diversification events reinforces the importance of considering the entire temporal framework of evolution within this group, as it reveals the contributions of different historical processes to their adaptation and speciation in the Eremaean Zone.

### The Pilbara as a biodiversity hotspot and cradle of novel adaptations

Our results clearly demonstrate that *N.* section *Suaveolentes* first occupied the more mesic portions of the Australian continent 6–5 Mya and then diversified (Fig. 5) in the Pilbara region. Aridity increases from the forested areas of the south-western and eastern coasts (mean annual rainfall of 1000–2500+ mm) to the hummock grasslands in the desert inland regions of the continent (< 200 mm; Groves, 1999). Mountains in the Pilbara are located near the ocean, providing a thermal buffer that maintains relatively stable temperatures. This proximity to the ocean and varied topography contributes to the region’s unique biodiversity and ecological stability. The plants and animals there are exceptionally diverse for an arid region (Booth *et al*., 2022), undoubtedly due to the complexity of the landscape with diverse soil types and topography, long-term geological stability, and presence of abundant refugia (e.g., multiple gorges with permanent water) providing mesic havens during even the most arid years (e.g., El Niño years). The high mountains of the Archaean Pilbara Craton (Hammersley Ranges at 1,500 m) differ from those of the Proterozoic Gascoyne Complex to the south, which are lower and the product of the collision of the Pilbara and Proterozoic Yilgarn Cratons (Myers, 1993), and the overall effect of this geological history on the Pilbara region has been the production of a mosaic of arid and mesic habitats in close proximity, making them unique on a planetary scale and a laboratory for diversification and evolutionary novelty.

The once tropical, forested center of Australia was long ago replaced by extensive arid lands beginning in the Miocene (23.03–5.33 Mya) reaching into the Pliocene (5.33 to 2.58 Mya) and Pleistocene (2.58 mya to 11,700 kya; Flower & Kennett, 1994). The arid center of the Australian continent has received much less attention (Byrne *et al*., 2008) than other regions, and most detailed studies of evolutionary diversity in Australia have focused primarily on the tropical rainforests (Bell *et al*., 2010) and temperate forests (Chapple *et al*., 2011; Kay & Keogh, 2012).

Some plant phylogeographic studies have focused on genetic diversity in the Pilbara bioregion, but most have examined single or pairs of species endemic to the Pilbara (Levy *et al*., 2016; Nistelberger *et al.,* 2020; Millar *et al*., 2022), in which they found high levels of variation that was not geographically structured due to gene flow across the region. The Pilbara bioregion has often been found to harbor high levels of diversity also at the species level, making it a biodiversity hotspot for both plants (Anderson *et al*., 2016) and animals (see below), but how much this species diversity has influenced broader biotic patterns in the EZ has been understudied. Comprehensive environmental surveys in advance of mining projects, 2002–2007 (McKenzie *et al*., 2009), resulted in the discovery of hundreds of new plant and animal species. For beetles and scorpions, 68% and 83%, respectively, could not be assigned to described species (Guthrie *et al*., 2010; Volschenk *et al*., 2010). Fine-scale genetic studies have suggested substantial cryptic diversity and complex genetic patterns across the Pilbara (Pepper *et al*., 2008, 2011a; Shoo *et al*., 2008; Doughty *et al*., 2010, 2011b; Catullo *et al*., 2011; Anderson *et al*., 2016). In contrast to our results, Ladiges *et al*. (2011) found species diversity in *N*. section *Suaveolentes* to be greatest in central Australia, where several of their “tracks” overlap, but our findings with highly revised species circumscriptions identify the Pilbara region as the most diverse in both species numbers and lineages.

The only other studies comparable in terms of nearly complete species-level sampling to ours are those of *Ptilotus* (Amaranthaceae; Hammer *et al*., 2021) and *Triodia* (Toon *et al.,* 2015; Anderson *et al*. 2016), which both found that the detected high levels of species diversity in the Pilbara could act as a source of pre-adapted xeric diversity for the rest of the EZ. However, it is important to note that the species-level sampling in Triodia was underestimated and included many incorrect identifications, making some interpretations problematic. Unlike *Ptilotus* and *Triodia*, which both arrived from Africa and Asia pre-adapted to aridity, the species of *N*. section *Suaveolentes* developed novel adaptation(s) to aridity *de novo* in the Pilbara before dispersing throughout the EZ. We suspect that a “key” adaptation for *N*. section *Suaveolentes* is strict inhibition of germination until the precise conditions for growth and seed production occur. They exhibit no succulence or other obvious physical attributes typically associated with aridity, and they disappear into the soil seed bank before the annual onset of summer drought and extreme heat, a key adaptation to surviving aridity. The species of *N*. section *Suaveolentes* thus experience the arid zone when it is neither too dry nor too hot, indicating that their distinct ancestral habitats (like those of tomato and potato), while different from current arid conditions, did not hinder their ability to find suitable environments for survival and subsequent conquest of the EZ.

Because of our improved species circumscriptions (not the different geographical areas underpinning their analysis), the main distinction between our conclusions and those of Ladiges *et al*. (2011) is that dispersal rather than vicariance is more explanatory in the evolution of *N.* section *Suaveolentes*. In Ladiges *et al*. (2011), dispersal only came into the picture as the explanation for species with broad distributions, which occurred after vicariance laid down the general patterns, whereas we envisage lineage diversification to have taken place in the Pilbara region, after which dispersal took place twice to other parts of the tropical/subtropical zones (i.e., the *N. benthamiana* and *N. occidentalis* clades; the former inhabiting only mesic sites), followed by the central and southern districts (the *N. simulans* clade) and finally the southern central and far southern areas (the *N. suaveolens* clade). In each of these clades, further diversification outside the Pilbara Region produced species inhabiting both mesic and xeric habitats, and to adequately address in which direction preferences changed more study of species limits/phylogenetics is needed. Most widespread species, e. g., the N. benthamiana complex (Cauz-Santos et al., 2022; Chase et al., 2022), appear to be species complexes, and we are adding more accessions to our analyses to try to address this issue. Dispersal-mediated founder events and genetic drift could have limited their evolutionary potential as the individual clades left the Pilbara, reducing their evolutionary potential and constraining further change, much as these new species were perhaps spatially constrained by their specializations. The data produced for this study cannot address these issues, but we have collected additional data for more accessions that will permit these topics to be investigated in future studies.

### The importance of evaluating species delimitation

For robust understanding of evolutionary history, species limits must be well understood. If species limits have not been assessed properly, then the distributions of such taxa are meaningless biologically. The previous phylogeographic analysis of *N*. section *Suaveolentes* (Ladiges *et al*., 2011) relied on the species as delimited by Horton (1981), which was the basis for the treatment in the *Flora of Australia* (Purdie *et al*., 1982), plus four others described after the *Flora* treatment. Here, we updated the analyses using a set of species with highly modified circumscriptions, including many newly recognized species (Chase *et al*., 2018, 2021, 2023). For example, in the wider Pilbara region (north-western Western Australia), only three of 15 species were recognized by Horton (1981); the remaining ten species have been described in the last five years, mostly discovered during recent fieldwork (Chase et al., 2021, 2023).

### Implications for conservation planning

Our results highlight the evolutionary significance of the Pilbara region, providing a foundation for prioritizing areas for conservation and developing management plans. The phylogenetic and biogeographical results presented in this study can inform conservation strategies by pointing areas with high species diversity and endemism, suggesting these as high-priority zones for conservation efforts. Protecting these areas can help preserve the evolutionary potential and ecological functions of the Pilbara’s unique flora.

Here, we show that the Pilbara region represents a major cradle of diversification in *Nicotiana*, which is echoed in other plant (Anderson *et al*., 2016) and animal groups (Ashman *et al*., 2018). It is an extensive, geologically complex, culturally important region for which only 6% sits in formally protected reserves (Government of Western Australia, 2017). However, the explosion of mining activity throughout the region in the past 40 years, with major mine expansions underway and planned, combined with an emerging knowledge of the biodiversity of the region highlight the risks of continued development in the absence of robust, detailed biological surveys by specialists in each group.

An alarming discovery from our study concerns the number of evolutionarily distinct *Nicotiana* species that appear to have extremely restricted distributions, particularly in the Hamersley Basin, the region comprising the unique iron-rich rocks at the core of the Australian mining industry. Extensive biotic surveys prior to mining notwithstanding (McKenzie *et al*., 2009), none of the new species of *Nicotiana* section *Suaveolentes* in the Pilbara region was identified, highlighting the need for detailed genetic and taxonomic studies as part of specialist treatments. This parallels the findings in Anderson *et al*. (2016) in *Triodia*, in which multiple new species were identified in the Pilbara (Barrett & Trudgeon, 2018; Barrett, 2019; Barrett *et al*., 2023). One of our new, narrowly distributed, cryptic-species discoveries from the Pilbara Craton is *Nicotiana karijini* (Chase & Christenhusz, 2018; previously identified as the more widespread *N. umbractica*), for which six of seven known collections were made during mine-site surveys. The possible fate of such species is obvious and leads us to speculate that many new species have gone extinct before they were described.

## Conclusions

Our results here have demonstrated that the large arid portions of Australia can act as a catalyst for rapid adaptation and diversification. While this phenomenon has been documented in some reptile groups, particularly arid-zone geckos (Pepper *et al*., 2011b; Pepper *et al*., 2013; Ashman *et al*., 2018), our study contributes providing detailed insights into plant diversification. Without first examining species limits, our biogeographic conclusions could never have been reached, and thus the first period of our studies focused on their taxonomy. The Australian arid zone is a relatively geologically stable and still largely undisturbed set of environments (despite numerous mines in areas like the Pilbara), making it an ideal setting for studies of speciation and diversification. Topographical heterogeneity combined with nearby marine influences have created localized regions within the Pilbara region with more buffered environments compared to the much larger and more homogeneous surrounding arid zone (Macphail & Stone, 2004; Byrne et al., 2008), permitting ancestrally mesic-adapted taxa like *Nicotiana* to experiment repeatedly with adaptations that have then allowed them to exploit available niches in the rest of the arid zone. Detailed molecular studies across the EZ are at present highly limited, and the high species diversity of the Pilbara Region, especially of cryptic, undescribed species attributable to its status as a mesic refuge, means that it should be an important focus of attention in the future.

Plants and perhaps also terrestrial invertebrates typically have more direct ties to the physical environment than vertebrates because they are inherently less vagile and thus more likely to provide important models to investigate genetic patterns/barriers across the EZ. The Pilbara is an ancient, topographically complex landscape of plateaus, gorges, valleys, and ranges with meteorological extremes and seasonal monsoons/cyclones. We hypothesize that the ancestors of the Pilbara lineages in *N.* section *Suaveolentes* entered the mesic refuges of the Pilbara Craton from the more coastally focused monsoonal region roughly 5 Mya. They became locally adapted to these various ancient and highly stable terrain types and subsequently were exposed repeatedly and became adapted to the interdigitated arid micro-habitats, which then permitted dispersals in several waves to other parts of the arid zone, including most recently (in the last million years) to sandy dune fields and the most homogenous, flat and extremely arid, southern parts of continent (e.g., the Nullarbor). The biotic history of this most ancient landscape, the Pilbara Craton, remains largely unknown and speculative. Our results should provide impetus to develop further an understanding of it and its contribution to the EZ flora and fauna.

## Supporting information

with Treeannotator v.1.8.3 (Drummond et al., 2012) (Fig. S1)

distribution of this section in central and southern Australia (Table S4, Fig. S2).

Sampling of 273 accessions from 58 species of N. section Suaveolentes (Table S1)

According to the Akaike information criterion (AIC), the best-fit model was DIVA+J (Table S2).

The BioGeoBEARS model testing for this distribution also resulted in the DIVA+J as best fit AIC model (Table S3).

leading to the current distribution of this section in central and southern Australia (Table S4, Fig. S2).

## Acknowledgements

We thank Juliane Baar and Daniela Paun for constructing RADseq libraries. Financial support to prepare the RAD library and analyze data were funded by research grants P26548-B22, P33028-B of the Austrian Science Fund (FWF) and an award (Dr. Anton Oelzelt-Newin Foundation) from the Austrian Academy of Sciences (ÖAW) to Rosabelle Samuel. The work was also supported by the Marie Curie individual fellowship for CondensDrought project (Grant agreement: 101029312) awarded to Luiz Cauz-Santos. Leaf samples of *N. fatuhivensis* were provided by Kenneth Wood, National Tropical Botanical Garden, Hawai’i. We thank Aureliano Bombarely for giving us earlier access to the *N. benthamiana* v.2.6.1 reference genome.

## Data availability statement

The data have been deposited in the NCBI Sequence Read Archive (BioProject ID PRJNA681916), and the files are available under the SRA Study SRP295424.

## Benefit sharing statement

Benefits from this research accrue from sharing our datasets on public databases as described above. Research collaborations were developed with scientists from the countries providing genetic samples, and all collaborators are included as co-authors.

## Author contributions

MWC and RS conceived this study. The fieldwork and collected/grew of accessions were conducted by MWC and MJMC. LACS, DM and OP analyzed the data. LACS and MWC wrote the manuscript with input from RS, KWD, JGC and OP.

