## Supplementary material for "Recent speciation and adaptation to aridity in the ecologically diverse Pilbara region of Australia enabled the native tobaccos (*Nicotiana*; Solanaceae) to colonize all Australian deserts": with Treeannotator v.1.8.3 (Drummond et al., 2012) (Fig. S1)

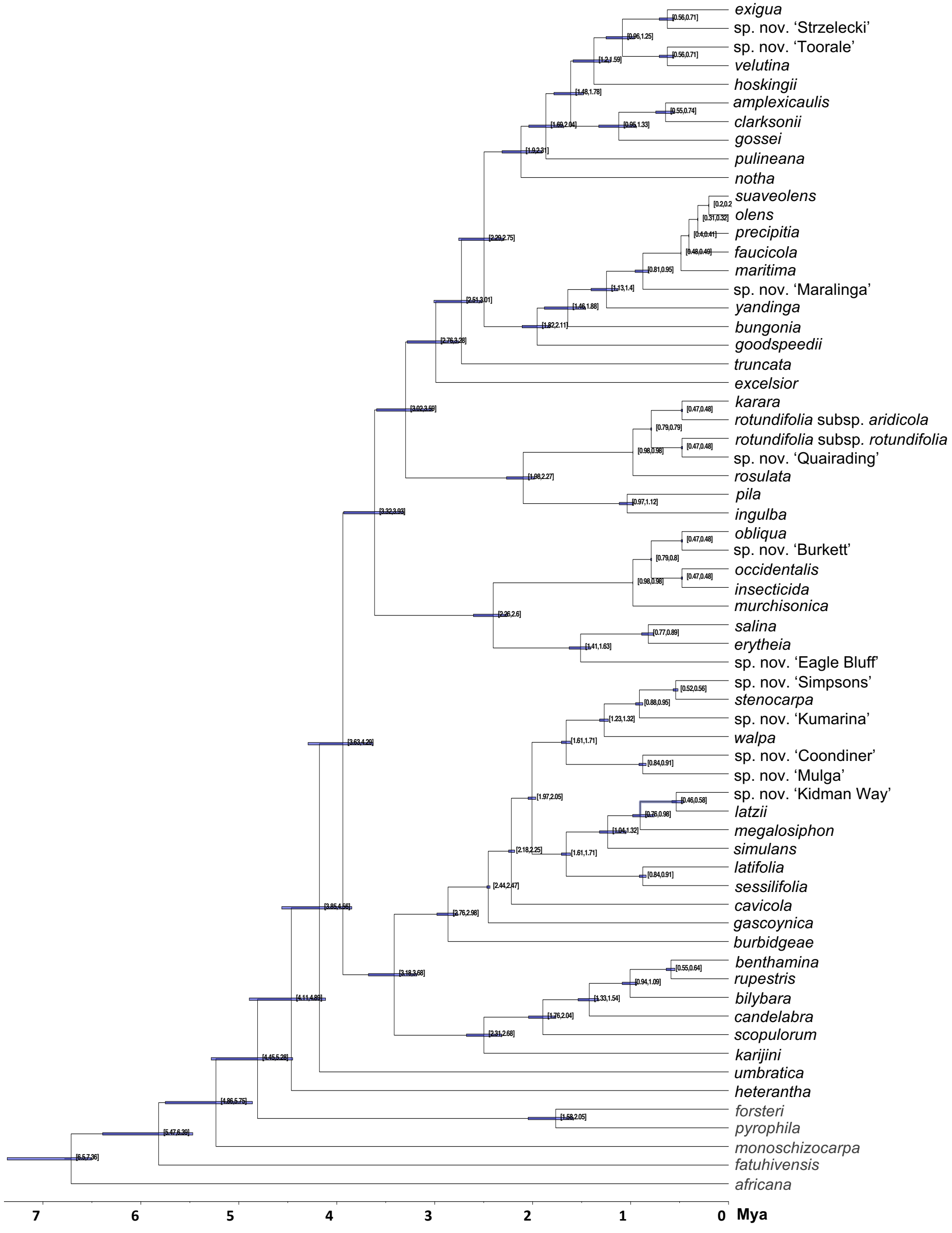


Fig. S1. Maximum clade credibility (MCC) tree derived from the bootstrap trees obtained in RAxML and dated in TreePL. The blue bars indicates the confidence intervals for the divergence time estimates.
