## Supplementary material for "Recent speciation and adaptation to aridity in the ecologically diverse Pilbara region of Australia enabled the native tobaccos (*Nicotiana*; Solanaceae) to colonize all Australian deserts": distribution of this section in central and southern Australia (Table S4, Fig. S2).

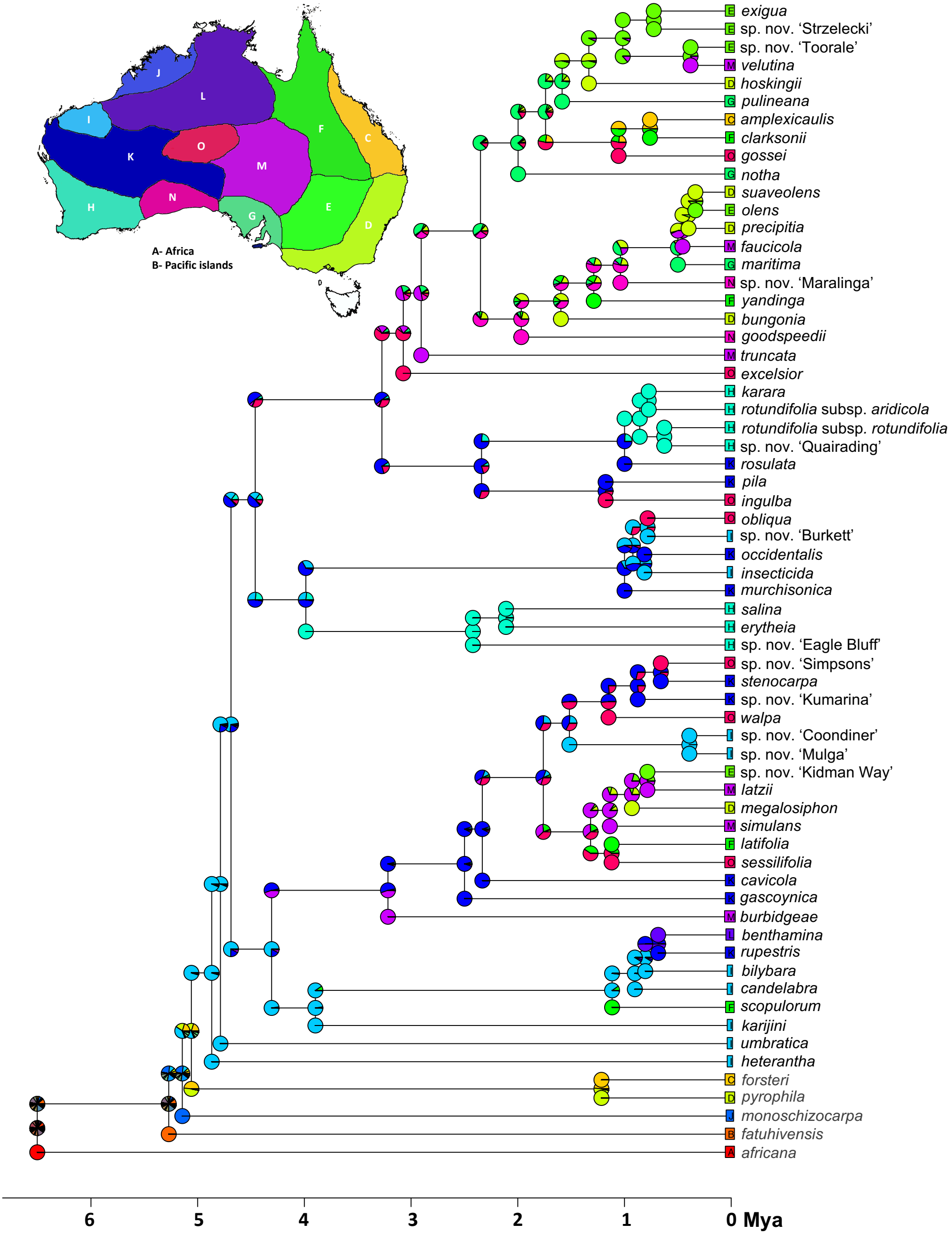


Fig. S2. Biogeographic history of *N*. section *Suaveolentes*. Ancestral area reconstruction using the operational areas adapted from Ladiges *et al*. (2011) and presented here in the map in the upper part of the figure.
