## Supplementary material for "Recent speciation and adaptation to aridity in the ecologically diverse Pilbara region of Australia enabled the native tobaccos (*Nicotiana*; Solanaceae) to colonize all Australian deserts": According to the Akaike information criterion (AIC), the best-fit model was DIVA+J (Table S2).

Table S2. The BioGeoBEARS model testing for the drainage divisions of Australia as operational areas: log-likelihood (-LnL); rate of dispersal (d) rate of extinction (e); rate of jump dispersal (j); Akaike information criterion (AIC), Akaike information criterion weighted (AIC_wt). Selected model is highlighted in red.

| Model | LnL | n* | d | e | j | AICc | AICc_wt |
| --- | --- | --- | --- | --- | --- | --- | --- |
| DEC | -225 | 2 | 0.025 | 0.17 | 0 | 453.9 | 8.4e-34 |
| DEC+J | -149.5 | 3 | 1.0e-12 | 1.0e-12 | 0.063 | 304.9 | 0.19 |
| DIVALIKE | -213.9 | 2 | 0.023 | 0.076 | 0 | 431.7 | 5.6e-29 |
| DIVALIKE+J | -148 | 3 | 1.0e-12 | 1.0e-12 | 0.061 | 302.1 | 0.79 |
| BAYAREALIKE | -247.8 | 2 | 0.021 | 0.35 | 0 | 499.6 | 1.0e-43 |
| BAYAREALIKE+J | -151.8 | 3 | 1.0e-12 | 1.0e-12 | 0.064 | 309.5 | 0.019 |

*Number of parameters
