## Supplementary material for "Recent speciation and adaptation to aridity in the ecologically diverse Pilbara region of Australia enabled the native tobaccos (*Nicotiana*; Solanaceae) to colonize all Australian deserts": The BioGeoBEARS model testing for this distribution also resulted in the DIVA+J as best fit AIC model (Table S3).

Table S3. The BioGeoBEARS model testing for the distribution areas adapted from Ladiges *et al*. (2011): log-likelihood (-LnL); rate of dispersal (d) rate of extinction (e); rate of jump dispersal (j); Akaike information criterion (AIC), Akaike information criterion weighted (AIC_wt). Selected model is highlighted in red.

| Model | LnL | n* | d | e | j | AICc | AICc_wt |
| --- | --- | --- | --- | --- | --- | --- | --- |
| DEC | -234.7 | 2 | 0.018 | 0.092 | 0 | 473.7 | 6.7e-35 |
| DEC+J | -156.6 | 3 | 1.0e-12 | 1.0e-12 | 0.074 | 319.7 | 0.19 |
| DIVALIKE | -231.8 | 2 | 0.021 | 0.050 | 0 | 467.8 | 1.3e-33 |
| DIVALIKE+J | -155.2 | 3 | 1.0e-12 | 1.0e-12 | 0.070 | 316.8 | 0.80 |
| BAYAREALIKE | -252.1 | 2 | 0.021 | 0.57 | 0 | 508.5 | 1.9e-42 |
| BAYAREALIKE+J | -159.4 | 3 | 1.0e-12 | 1.0e-12 | 0.076 | 325.3 | 0.011 |

*Number of parameters
