## Supplementary material for "Recent speciation and adaptation to aridity in the ecologically diverse Pilbara region of Australia enabled the native tobaccos (*Nicotiana*; Solanaceae) to colonize all Australian deserts": leading to the current distribution of this section in central and southern Australia (Table S4, Fig. S2).

Table S4. Biogeographic stochastic mapping (BSM) counts for *Nic*. sect. *Suaveolentes* using the operational areas adapted from Ladiges *et al*. (2011) and the DIVALIKE+J model selected in BioGeoBEARS. Mean values (mean) and standard deviations (stdev) are event counts of 100 BSMs.

| Mode | Type | mean | stdev | % |
| --- | --- | --- | --- | --- |
| Within area speciation | Sympatry - narrow | 19.45 | 1.42 | 30.87 |
| Dispersal | Jump-dispersal | 41.2 | 1.52 | 65.39 |
| Vicariance |  | 2.35 | 0.77 | 3.73 |
| Total events |  | 63 |  |  |
